# RhizoVision Explorer: Open-source software for root image analysis and measurement standardization

**DOI:** 10.1101/2021.04.11.439359

**Authors:** Anand Seethepalli, Kundan Dhakal, Marcus Griffiths, Haichao Guo, Gregoire T. Freschet, Larry M. York

## Abstract

Roots are central to the function of natural and agricultural ecosystems by driving plant acquisition of soil resources and influencing the carbon cycle. Root characteristics like length, diameter, and volume are critical to measure to understand plant and soil functions. RhizoVision Explorer is an open-source software designed to enable researchers interested in roots by providing an easy-to-use interface, fast image processing, and reliable measurements. The default broken roots mode is intended for roots sampled from pots or soil cores, washed, and typically scanned on a flatbed scanner, and provides measurements like length, diameter, and volume. The optional whole root mode for complete root systems or root crowns provides additional measurements such as angles, root depth, and convex hull. Both modes support providing measurements grouped by defined diameter ranges, the inclusion of multiple regions of interest, and batch analysis. RhizoVision Explorer was successfully validated against ground truth data using a novel copper wire image set. In comparison, the current reference software, the commercial WinRhizo™, drastically underestimated volume when wires of different diameters were in the same image. Additionally, measurements were compared with WinRhizo™ and IJ_Rhizo using a simulated root image set, showing general agreement in software measurements, except for root volume. Finally, scanned root image sets acquired in different labs for the crop, herbaceous, and tree species were used to compare results from RhizoVision Explorer with WinRhizo™. The two software showed general agreement, except that WinRhizo™ substantially underestimated root volume relative to RhizoVision Explorer. In the current context of rapidly growing interest in root science, RhizoVision Explorer intends to become a reference software, improve the overall accuracy and replicability of root trait measurements, and provide a foundation for collaborative improvement and reliable access to all.

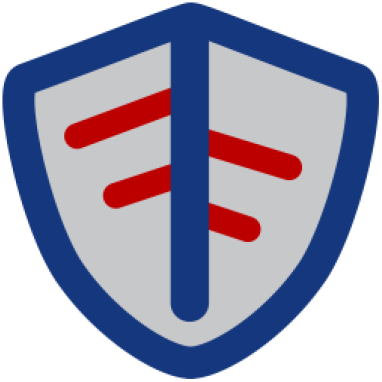

## Introduction

Roots are central to understanding and providing answers to pressing global challenges like climate change’s impact on plant communities, ecosystem services, and food security. From an ecological perspective, roots are an integral component of global biogeochemical processes, accounting for 46% of the net primary productivity of the terrestrial biosphere globally, ranging from 32% in croplands, 41%–58% in various forest types, and 64% in grasslands (Gherardi and Sala, 2020). They have demonstrated influence on countless numbers of plant and ecosystem functions (Bardgett et al. 2014; Freschet et al. 2020). As such, on an ecosystem level, conceptual understanding, parameterization, and spatial scaling of various root processes are critical for quantifying carbon, water, and nutrient fluxes (Warren et al. 2015). From an agricultural perspective, rising demand for food, increased cost of production, agricultural intensification, and depletion of finite resources in the face of changing climates have adverse effects on food security prospects (Godfray et al. 2010). Crops with superior root phenotypes are considered an important determinant of future food security, and the key to the second green revolution that can improve farm productivity and sustainability (Lynch 2007). Additionally, the root-soil rhizosphere is regarded as the keyspace for the future discovery of novel root traits relevant to increasing crop and farm productivity (Tracy et al. 2020). Hence, a better understanding and characterization of belowground plant components will enable a greater knowledge of the underlying ecosystem processes in both natural and agricultural systems.

Due to their vital roles in anchorage, acquisition of water and nutrients, and driving soil biology, plant root systems have been studied by the scientific community for over 100 years (Lux and Rost, 2012). Roots still remain the least understood plant organ, partly because their underground growth is obscured by the opaque soil matrix (Eshel and Beeckman 2013; Hochholdinger 2016). This belowground life leads to the need to sample roots by excavation or soil coring, followed by washing, which requires substantial effort. However, the rewards for measuring roots are great, with substantial evidence presented for how numerous root traits directly influence processes like nitrogen uptake and soil reinforcement (Freschet *et al*. 2020). The term “phenotyping” has become commonly used to mean measuring traits in the context of screening crops for genetic mapping. Typically, these applications require measuring hundreds or thousands of samples, and so high-throughput phenotyping is the goal (Fiorani and Schurr 2013). In this manuscript, we assume that the need for reliable, fast, and precise phenotyping is needed for research in both agricultural and natural systems.

From a historical perspective, some of the primary root measurements used have included root dry weight, number, volume, surface area, and length. As explained by (Böhm 1979), the dry weight is commonly used due to its relative ease and, when measured on distinct organs of a plant, is a good indicator of carbon allocation in the plant. Volume is known to correlate with root dry weight in many circumstances and is relatively easy to measure by water displacement methods (Harrington *et al*. 1994; Pang *et al*. 2011). Although root mass or volume measurements are relatively easy to measure, they cannot adequately explain many root functions in the plant-soil continuum (Costa *et al*. 2000). Böhm (1979) explained that volume did not become widely used because these parameters cannot distinguish samples with few thick roots from those with many fine roots, nor can dry weight. By 1970, Gardner (1964) and others had shown that root length was the best predictor of water uptake, and so was highly desirable to measure. Throughout the 20th century, root length was typically measured by placing roots on graph paper, using tacks or gum to hold the ends in place to stretch out, and lateral roots had to be cut away to measure separately (Böhm 1979). Therefore, any method for measuring the length and other parameters with higher throughput and precision was highly sought.

Methods designed to overcome the barrier in measuring root length using a ruler or graph paper date back at least to Newman’s (1966) “line-intersection” method, assuming random placement of lines and its variant by Tennant using a grid of lines (1975). In these methods, total root length is estimated by manual counting of roots that intersect a line marker and a formula to convert to root length. Subsequently, the line-intersection method was digitized using a photo-electric sensor that traveled over the top of a backlit root sample to count transitions from white to black using a scaler or electronic counter (Rowse and Phillips 1974). Image-based methods were employed to capture mini-rhizotron images starting in the 1970s, and these images were processed with the line-intersection method, either manually or with the digitized version (Upchurch and Ritchie, 1983). Smucker et al. (1987) described a complete algorithm for thresholding a root image, thinning to produce a skeleton, using erosion to compute a distance map, and calculating length, surface area, and volume from images acquired using a videotape camera. In parallel, more advanced digitized methods for the line-intersection method were still in development in 1989 using both specialized counting devices as described above (Harris and Campbell, 1989), but also image analysis to count from images acquired by a desktop document scanner (Krstansky and Henderson 1989). The business applications of desktop scanners led to their wider availability and lower costs, while the optical arrangement made the method superior relative to using cameras; leading to higher resolution images with less distortion. The first reported use of a desktop scanner for acquiring images of spread roots followed by image analysis of the entire root objects (rather than line-intersection) was described by Pan and Bolton (1991), with the algorithm determining the perimeter of root objects and then deriving root length, area, and average diameter from that single measurement. WinRhizo™ is a commercial and closed-source software released in 1993 (Arsenault *et al*. 1995) based on the principle of standardizing the use of desktop scanners and image analysis using similar algorithms as described by Smucker et al. (1987) to measure root length, diameters, areas, and volumes. ROOTEDGE is free and open-source software developed to measure root length from scanned images using the edge-chord algorithm (Kaspar and Ewing 1997), for which a legacy DOS program was still available for download as of the writing of this article (https://www.ars.usda.gov/midwest-area/ames/nlae/docs/software-available-from-nlae/). As root image analysis methods focused on using skeletonization routines for measuring length, Kimura and Yamasaki (2003) provided a useful refinement of the algorithms to measure length and diameters more accurately, using the NIH Image software (now ImageJ).

Since then, advances in image capture and image processing algorithms have continued and been widely utilized to measure various root metrics. Roots are imaged across a wide degree of modalities, such as *in situ* with minirhizotrons or soil pits, in rhizoboxes, on colored backgrounds, or with flatbed scanners (Topp *et al*. 2016). Therefore, image analysis approaches are often specific to particular types of collected images. At the time of writing, there are 17 root image analysis tools for single root images and 42 tools for root system images listed on Quantitiative-Plant.org, each with varying features and assumptions for input image type (Lobet *et al*. 2013). At present, 2D root imaging and image analysis is the most popular root phenotyping approach, after root dry weight determination, as it is the most accessible and simplest image data to acquire and analyze. Software that work with 2D branched and connected root system images such as root crowns or seedling root systems grown on blue paper included EZ-Rhizo (Armengaud *et al*. 2009), SmartRoot (Lobet *et al*. 2011), RootNav (Pound *et al*. 2013), ARIA (Pace *et al*. 2014), DIRT (Das *et al*., 2015), and RhizoVision Analyzer (Seethepalli, Anand *et al*., 2020). In many cases, roots are not connected as they have been excavated from the field or pots, and these roots are typically imaged on a flatbed scanner then analyzed using the commercial WinRhizo™ software (Regent Instrument Inc., Quebec, Canada) or free software including IJ_Rhizo (Pierret *et al*. 2013); requiring ImageJ), GiA-Roots (Galkovskyi *et al*. 2012); no longer available for download), and saRIA (Narisetti *et al*. 2019; requiring Matlab runtime). While the growing numbers of image analysis tools for root phenotyping have been really useful to the root science community, these have also introduced a range of biases and inconsistencies among studies (e.g., Rose and Lobet 2019) and likely reinforced the conceptual barriers between root researcher communities (see Freschet *et al*. 2020). Currently, there is a need to bring these tools together into a user-friendly, generalist, all-inclusive package that has wide utility across plant science, provides standardized results, and is open-source.

As presented in detail below, RhizoVision Explorer fills the gap for generalized root image analysis, with modes to extract several root traits from both connected and unconnected plant root systems. Here we describe aspects of the user-friendly, lightweight, and open-source graphical user interface that facilitate high-throughput root phenotyping, such as its ease to install and use, support of multiple image formats, the inclusion of image pre-processing, and batch analysis mode. Additionally, we present the methods and outcomes of the software validation using a novel copper wire image set and simulated roots with ground truth measurements. For scanned roots, the software was further compared with WinRhizo™ using images from several plant species and functional categories.

## Materials and Methods

RhizoVision Explorer (Fig 1) builds upon the functionality and codebase of RhizoVision Analyzer (Seethepalli and York 2019) to analyze not only root crown images but also broken roots that are imaged on a flatbed scanner. The term ‘Explorer’ indicates the user has more freedom to understand the analysis process and work interactively to optimize the desired outcome. The program adds functionality such as greater interaction with the raw, segmented, or analyzed image, performing root pruning, and support for the region of interest (ROI). The minimum-tested system requirements for the RhizoVision Explorer are an Intel or AMD x86 64-bit processor and a minimum of 4 GB of RAM. If the processor supports Intel AVX 2.0, the program is accelerated using AVX 2 vector processing instructions. The program does not require any installation or external dependencies and runs locally on Windows 8.1 or higher, but could be compiled to run on other operating systems in the future. The program is open-source, and the zipped binary files for simple installation are available for download at https://doi.org/10.5281/zenodo.3747697 (Seethepalli and York 2020).

**Figure 1.**
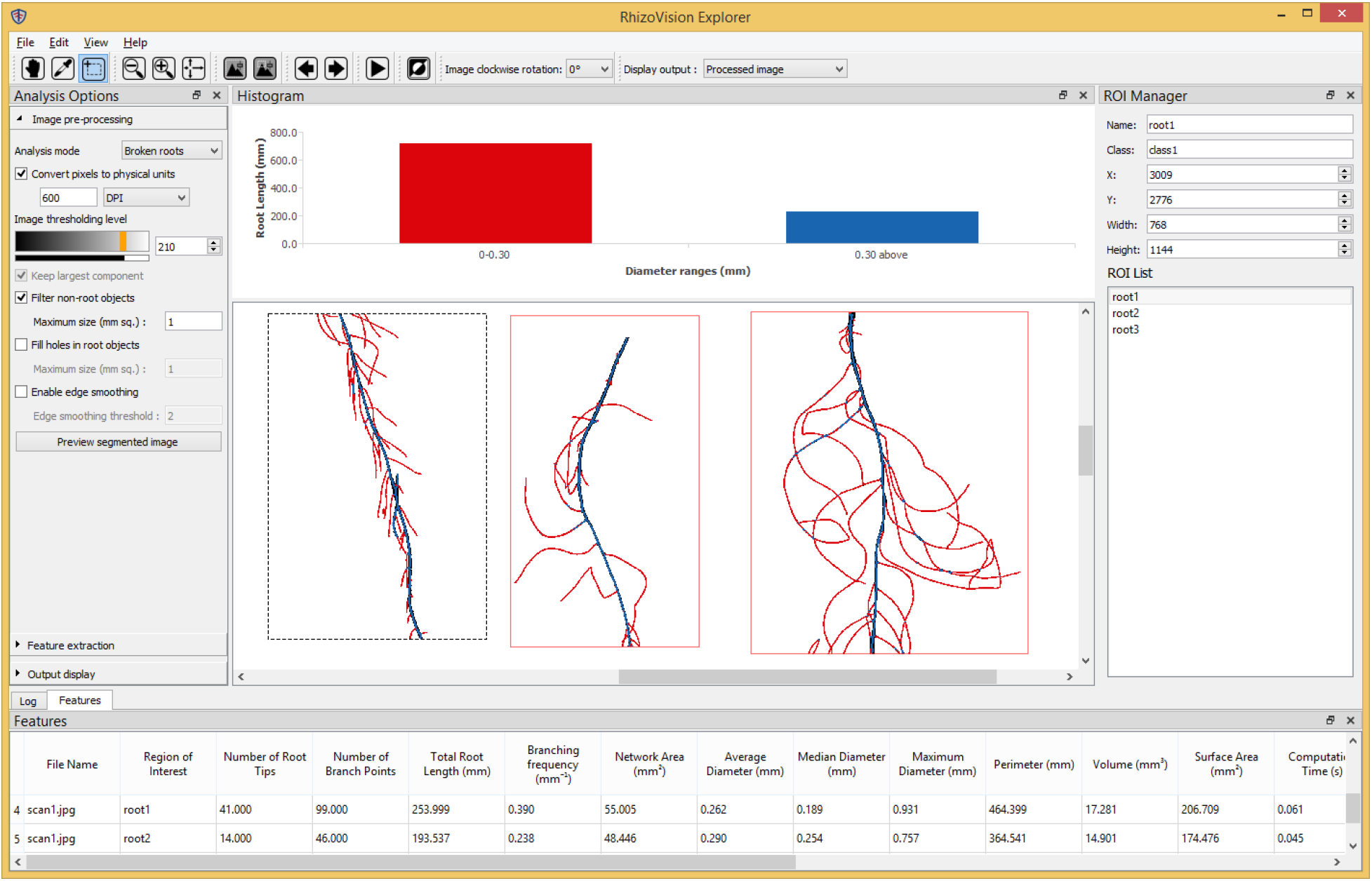
The RhizoVision Explorer application window consists of menus (top), toolbar under menus, Analysis Options pane (left), image window (center), Histogram pane (over image window), ROI Manager (right), and the Features or Log pane (bottom).

### Software Architecture

RhizoVision Explorer is written in C++ (https://github.com/noble-research-institute/RhizoVisionExplorer) and uses *cvutil* (https://github.com/noble-research-institute/cvutil), a C++ library that includes several helper functions that extend the *OpenCV* library (www.opencv.org) and was developed by the same team in parallel. These functions include determining the distance map, creating the skeletal structure, and smoothing the contours of a segmented image. Additionally, the *cvutil* library supports adding a user interface to an application with the main window, extending the user interface with the plugin system, and optionally enabling the ROI system that can be utilized by the plugins loaded by the application. Finally, the library includes several plotting functions for data visualization. The *cvutil* library utilizes the platform-independent *Qt* C++ library (www.qt.io) to generate the required graphical user interfaces. Using *OpenCV* and *QT* enables extending the software and required libraries to be deployed on operating systems other than Windows in the future.

The primary algorithms for extracting features, or measurements, from root images, were implemented within a plugin to the image processing window. The plugin also includes the logic for identifying the root topology and generating the feature image with visual elements such as the skeleton or convex hull overlaid on the segmented image. When the user starts analysis from RhizoVision Explorer, the main window checks if the image has any ROIs drawn by the user. If any ROIs are found, then the main window invokes the plugin for extracting features for every sub-image given by the ROI set. Otherwise, the whole image is passed to the plugin for analysis. The plugin then analyzes the image and returns the extracted features and the segmented and processed images to the main window. The main window then updates the features table with the extracted features, displays the processed image in the image pane, and updates the bar plot root length histogram that groups length measurements based on the diameter ranges given by the user before analysis.

The main window allows the user to change the analysis options interactively and run the analysis, thereby allowing the user to find the optimal analysis options for an image. These analysis options can be saved in a CSV file and loaded later. Similarly, the main window also supports loading and saving the ROIs as annotation text files, so that these annotations can be reused later.

### Description of analysis tools in RhizoVision Explorer

A typical workflow for analyzing a plant root image is to load the image in RhizoVision Explorer by drag-and-drop or through the file > open menu. Most common image formats are supported, including PNG, JPEG, BMP, and TIFF, in both grayscale and color, but note only the red channel is used for color images. The Analysis Options pane on the left side of the program lists the options for image preprocessing, feature extraction, and how the output of the analyzed image is displayed to the user. The user needs to set the options to be appropriate for any given experiment, preview segmentation, and then run the analysis. Upon pressing the ‘run analysis’ arrow button, the program extracts relevant features from the plant root image, displaying the processed image at the center of the window. The default analysis mode is for broken roots, but the software also has a whole root mode that will perform an analysis similar to RhizoVision Analyzer including extra features such as the convex hull, root system width and depth, and angle measurements. The user may convert pixels to physical dimensions in units of either pixels per mm or DPI (dots per inch) as typically used when scanning. Apart from the thresholding and color inversion options, which are present in the RhizoVision Analyzer, the program also supports filtering from the background (non-root objects) and the plant root portions (hole filling). Non-root object filtering enables soil particles or other debris, which may be present after root washing, to be filtered from the image. Hole filling is useful in cases where the root texture in the image may contain bright spots that may be identified as background after thresholding, and allows the small areas to be filled in that are surrounded by root pixels. The filtering is performed based on the user-provided maximum size of a filtered component for the background and the root portions of the image. The maximum size is provided in sq. mm if pixel to millimeter conversion is specified in the program. The program then filters the connected components based on the size of each component. Fig 2 shows the thresholded images of a root image patch before and after filtering noisy components.

**Figure 2.**
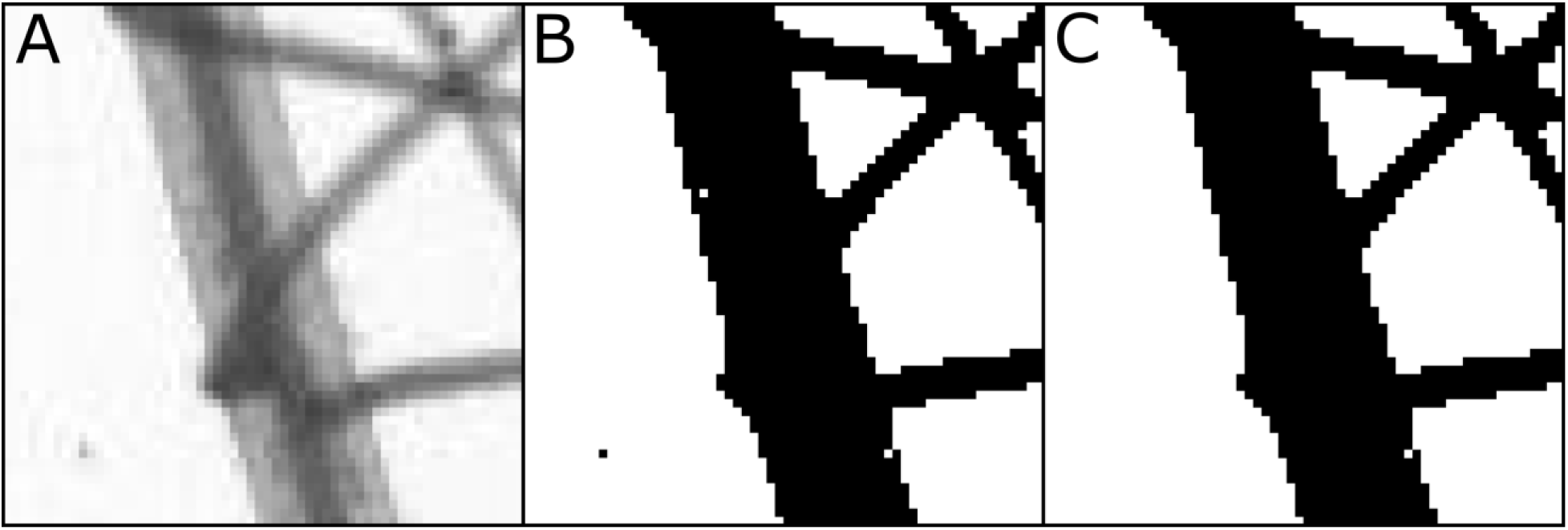
(A) Selected region of a plant root image taken from a flatbed scanner with a transparency unit. (B) Segmented image without noise filtering. We observe that the dark component on the white background is caused due to soil particles and a white component on the root portion is caused due to brighter root texture. (C) Segmented image with noise filtering. The white pixel at the bottom is not considered as a hole because the pixel is only surrounded by seven root pixels, rather than eight.

The contours, or root surfaces, of the thresholded and filtered image, can be optionally simplified using the Ramer-Douglas-Peucker line simplifier (Ramer 1972; Douglas and Peucker 1973). The procedure simplifies the contours of the thresholded image within a user-provided pixel distance threshold. Small contours of the segmented root can lead to the formation of short lateral roots during skeletonization that are not valid. Therefore, smoothing the surface using line simplification can reduce the number of these unwanted invalid lateral roots. It is recommended not to set values higher than 2.0 for pixel distance threshold for line simplification, because it may alter the root topology, leading to an angular appearance.

Using the segmented image generated using the methods described above, a precise Euclidean distance transform (Felzenszwalb and Huttenlocher 2012) is computed for the image (Fig 3A). The distance transform at any pixel is the shortest distance from the pixel to a background pixel. The skeletonization of the segmented image is performed by the identification of ridges of the distance transform (Fig 3B). This is done by checking each distance transform pixel if it is greater than two neighboring pixels of opposite directions. The ridges so identified are not always connected throughout the root system. This is because the distance transforms pixels at the junction of the main root and its lateral root increase towards the ridge of the main root. Hence, no ridge pixels can be found that can connect the lateral root to the main root. These ridges are connected in the steepest ascent algorithm to form a connected skeletal structure. The ridge may be 2 pixels wide in some places which may be formed in the ridge detection procedure. In such a case, the ridge is thinned using the Guo-Hall thinning algorithm (Lam *et al*. 1992) to one-pixel width while preserving the connectivity of the skeletal structure. The skeletal structure is internally stored as an image where pixel values take on the value computed from the distance map for that pixel and use it for computing the average diameter of all roots and to classify root segments into diameter bins as described below.

**Figure 3.**
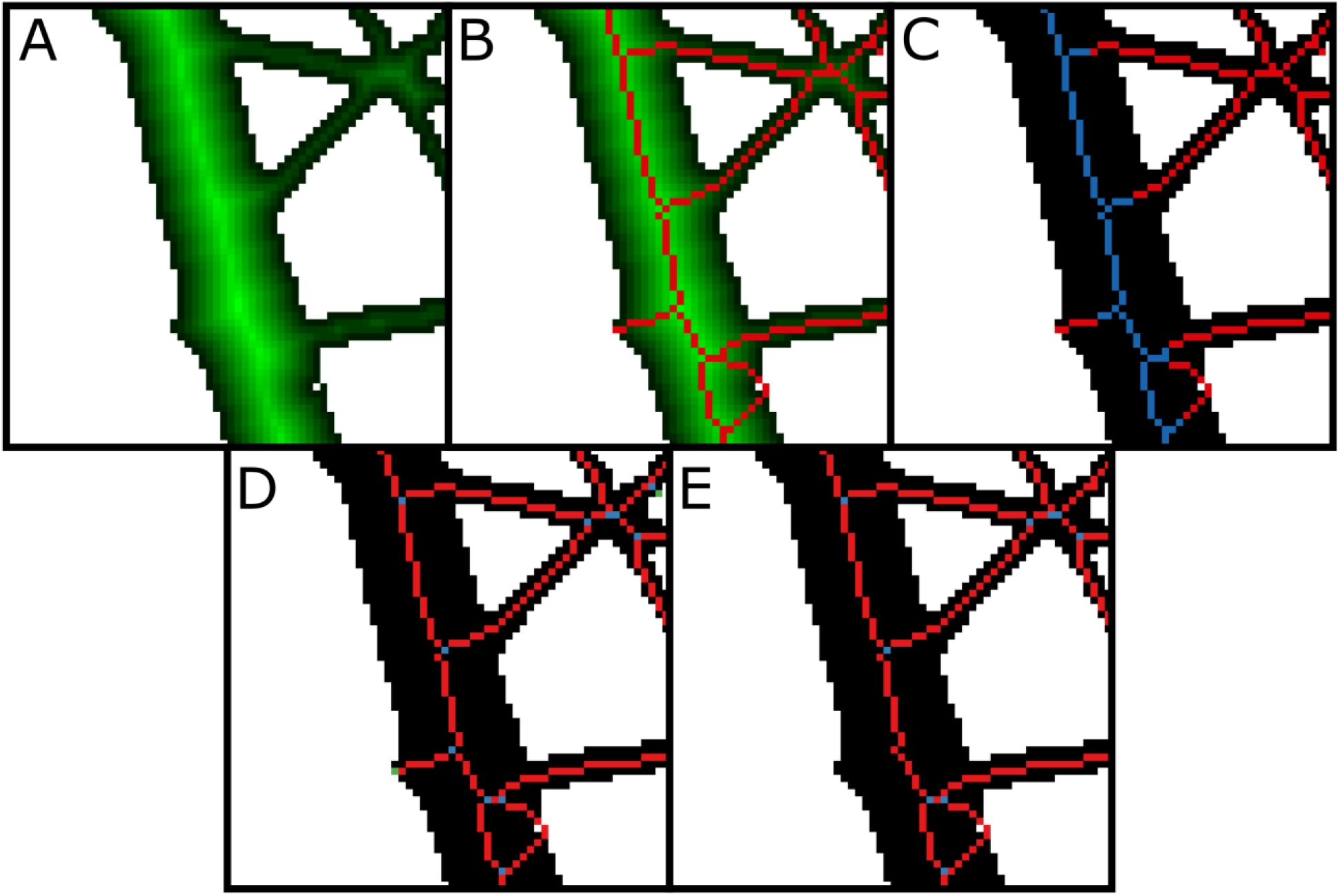
(A) Distance transform of the segmented image of the selected region shown in Fig 2A. (B) The skeletonization procedure finds the ridges and connects them using the steepest ascent algorithm. (C) Skeletal pixels colored based on the diameter ranges given in the feature extraction options. (D) Root topology without root pruning. The blue pixels are the branch points, and the green pixels are root tips. (E) Root topology with root pruning. Note two lateral roots in the bottom right of the image are not pruned because they form a single segment connecting two branch points.

After skeletonizing the segmented image, the topology of the root is identified, which involves finding the topological branch points, endpoints, and the root segments between any two branch points or a branch point and an endpoint (Fig 3D). The endpoints, here on called root tips, are identified as any skeletal pixel having only a single neighboring pixel and branch points have 3 or 4 neighboring pixels. This topology is stored as a copy of the skeletal image where pixels can take discrete values that encode the type, tip, branch point, or segment. Using this topological image and the skeletal image, another method for decreasing the number of short, invalid lateral roots was devised as an analysis option called “root pruning” as an alternative to using line simplification of the segmented image. The basic premise of root pruning is that the length of invalid lateral roots will be no longer than the radius of the parent root from which they branch. When the root pruning option is selected, the software evaluates the topological image to analyze all root segments starting from a branch point and ending in a tip. The software determines whether the length from the branch to the tip is longer than the radius of the branch point plus an additional number of pixels supplied by the user (Root pruning threshold). If a given lateral root is too short, then its tip and segment are deleted from the topological image and the skeletal image. If the resulting parent branch point has one neighbor, it becomes a tip; if two neighbors, it becomes merged with the segment to which it belongs; and if it still has three or more, it remains a branch point. Since new tips can emerge during this process, root pruning is iterative until no invalid lateral roots remain. Unlike line smoothing, root pruning does not alter the texture of the segmented root image. Root pruning removes the invalid lateral roots that can inaccurately affect the extracted features, such as the total root length (Fig 3E). The resulting pruned topological image is used for extracting features requiring lengths and diameters. Features extracted in RhizoVision Explorer that rely on the segmented image, the skeletal image, and the topological image are defined in Table 1. Surface area and volume are calculated assuming that every pixel in the skeleton is a cylinder, calculating based on its height and radius, and summing those for the entire image or diameter range.

**Table 1.** List of features extracted from a plant root image. Features extracted from broken roots only are marked with (*), and from whole root images only are marked with (**).

| Feature extracted | Description |
| --- | --- |
| Number of Root Tips | The number of root tips is pixels in identified root topology that have only one neighboring skeletal pixel. |
| Total Root Length (px, mm) | The sum of the Euclidean distances between the connected skeletal pixels in the entire root topology of the plant root image. |
| Network Area (px<sup>2</sup>, mm<sup>2</sup>) | The total number of pixels in the segmented image. |
| Average, Median, and Maximum Diameter (px, mm) | The distance transform value at each skeletal pixel is the radius at that pixel and is doubled to give the diameter. The average, median and maximum diameters are computed across all these skeleton pixels. |
| Perimeter (px, mm) | The sum of the Euclidean distances between the connected contour pixels in the entire segmented image of the plant root. |
| Volume (px<sup>3</sup>, mm<sup>3</sup>) and Surface Area (px<sup>2</sup>, mm<sup>2</sup>) | Using the radii for each skeletal pixel determined earlier, the volume at the skeletal pixel is calculated as the length of the pixel multiplied by the cross-sectional area of the root at that pixel. Similarly, the surface area is calculated as the length of the pixel multiplied by the circumference of the cross-section of the root at that pixel. Volume and surface area are calculated as the sum of values from all skeletal pixels. |
| Computation Time (s) | The time taken to analyze a plant root image is noted as computation time in seconds. |
| Root Length, Projected Area, Surface Area, and Volume histograms | For each skeletal pixel, the root length, projected area on the surface of the image plane, surface area, and volume of the root at that pixel are computed. These values are binned according to the diameter ranges specified by the user. |
| Number of Branch Points* | The number of branch points is pixels in the identified root topology that have at least three neighboring skeletal root segments. |
| Branching Frequency (px<sup>-1</sup>, mm<sup>-1</sup>)* | The number of branch points per unit root length. |
| Median and Maximum number of roots** | For each row in the segmented image, a horizontal line scanning is performed that records the number of pixel transitions from a background to a root pixel. This list of pixel transitions is sorted and the median and the maximum number of roots are noted. |
| Depth, Maximum Width (px, mm)** | The maximum vertical and horizontal distance the root crown grew at the time of imaging are noted as Depth and Maximum Width respectively. |
| Width-to-Depth Ratio** | The ratio of Maximum Width to Depth is noted as the Width-to-Depth ratio. |
| Convex Area (px<sup>2</sup>, mm<sup>2</sup>) and Solidity** | The area of the convex hull that is fit to include the entire root crown system is noted as Convex Area. Solidity is the ratio of the Network Area to Convex Area. |
| Lower Root Area (px<sup>2</sup>, mm<sup>2</sup>)** | For a root crown image, the skeletal pixel that has the maximum radius is located. The network area of the root system that is below the skeletal pixel located above is noted as Lower Root Area. |
| Holes and Average Hole Size (px<sup>2</sup>, mm<sup>2</sup>)** | Holes are background components between roots in a segmented image. These holes are counted, and their average size is determined. |
| Steep Angle Frequency, Medium Angle Frequency, Shallow Angle Frequency, and Average Root Orientation** | Within 40x40 pixel locality for every skeletal pixel in the center, we get the coordinates of all the skeletal pixels in that locality and compute the angular orientation for the locality. We group these orientations in bins of 0° to 30°, 30° to 60° and 60° to 90° and note the frequencies as Steep, Medium, and Shallow Angle Frequencies. The average of these orientations is noted as Average Root Orientation. |

Apart from extracting the root features, the program can also be used to visualize the analyzed root interactively. Further, after running the analysis, a histogram appears of root length grouped by different root diameter ranges provided by the user (Fig 1). Fig 3 shows different ways a processed root can be visualized. The visualization of a processed image contains options for displaying the distance transform map of the segmented image and generated skeletal structure, or medial axis. Additionally, for root crown images, the processed image can be optionally displayed with a convex hull that fits the whole root system, the holes, or the background components separated by the root system and contour of the segmented image. The skeletal structure can be displayed based on the user-specified diameter ranges (Fig 3C) or based on the topology of the skeletal structure (Fig 3D and Fig 3E). When a root skeletal structure is visualized based on diameter ranges, the ranges can be selected by the user to distinguish the first and second-order lateral roots (Fig 3C). This also changes the histogram features that are extracted, to get useful information such as the root length of the main roots and the lateral roots separately. When generating the skeletal structure based on root topology, the branch points and root tips are highlighted.

RhizoVision Explorer also supports several tools that dramatically improve the usability of the software. One such tool is the region of interest system, termed the ROI Manager. The ROI tool on the toolbar allows drawing multiple ROIs, and the ROI Manager allows custom naming of a selected ROI; manually changing XY coordinate position, width, and height; and maintains a list of ROIs in the image. For each ROI in an image, the roots within that ROI are analyzed separately and reported as a unique row of data, including a column for the name of the ROI. ROIs can be saved and loaded as an annotation file and are honored for batch processing.

### Validation datasets and procedures

For validation of data extracted from RhizoVision Explorer, known gauges of copper wire (Fig 4B) were used. Copper wire samples with average lengths of 30.48 cm and various diameter wire gauges (American Wire Gauge 40, 32, 28, 22, 16, and 10; with the diameter ranging from 0.07 mm – 2.58 mm) were used for the validation. For each gauge, two wires were used for a total of 12 wires. The diameter of each wire was measured with a digital micrometer and the length was measured by fitting string to the length of the wire, then measured with a ruler. The surface area and volume for each wire were calculated assuming the wires were cylinders. The average diameter for scans containing multiple wires was calculated as the weighted diameter based on the length of each wire in the scan. The wires were then scanned on an Epson Expression 12000XL flatbed scanner with a transparency unit (overhead light), as common practice for root imaging. The images were digitized at 600 dpi as 16-bit grayscale images and then saved as TIFF (tagged image file format). Scans were taken in 28 combinations from individual wires to all as shown in Table S1 in order to achieve variation in total length, average diameter, total surface area, and total volume. The copper wire image set used here is available in a public repository and can be downloaded at http://doi.org/10.5281/zenodo.4677546 (Dhakal *et al*. 2021a).

**Figure 4.**
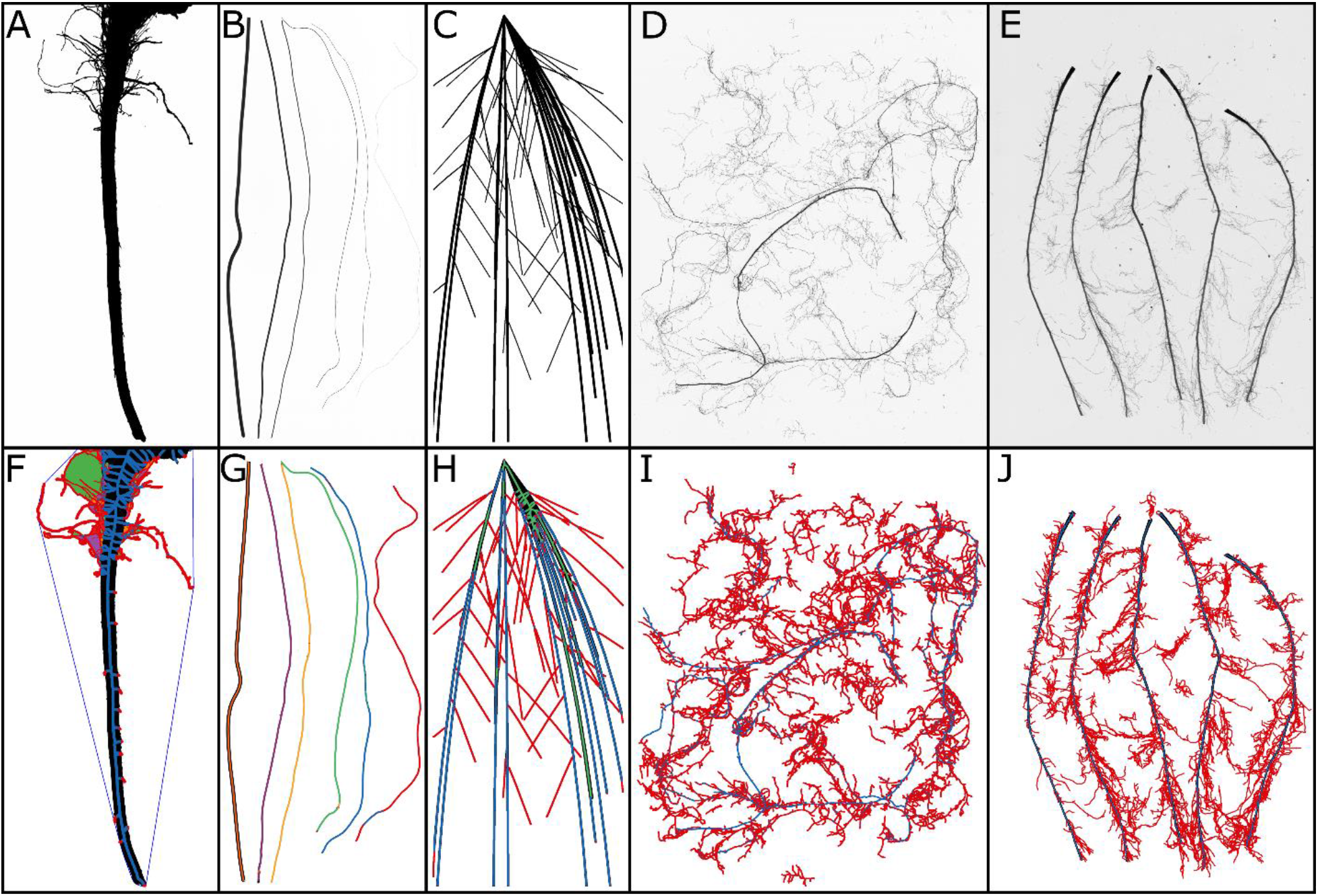
RhizoVision Explorer works with images that have a high contrast of roots with the background. (A) Whole root mode is intended to be used with connected root systems such as this soybean root crown image taken using the RhizoVision Crown platform. (B) A new image set containing copper wires of various diameters and combinations was created for validation. (C) Simulated root systems were analyzed from Rose and Lobet (2018). (D) Complex root samples in broken root mode such as from the herbaceous species dataset can be accommodated. (E) Maize nodal roots exhibit a substantial difference in diameter between nodal roots and their lateral roots. F, G, H, I and J depict corresponding feature images highlighting the additional measures in whole root mode in F and the ability to use diameter bins to accurately classify wires and roots (G, H, I, and J).

The wire images were processed in RhizoVision Explorer v2.0.2 (Fig 4G) using the following analysis options: Root type was chosen as broken roots, image thresholding level was set at 191, filter noisy components on the background was set as true, with the maximum noisy component size of 8 mm^2^. The threshold value was adjusted until the copper wires were fully visual, without introducing any deviation in the shape of the wires. The root pruning threshold was set to five pixels. Images were then processed in batch mode. The same images were processed in WinRhizo™ v2019a using the following analysis options: root detection was based on gray level and thresholding was set manually at 191 to match the options chosen for RhizoVision Explorer (RVE). Roots were chosen as darker than the background. Debris and rough edges removal were selected as low.

For comparison of performance metrics of RVE, we chose two popular software for root phenotyping, WinRhizo™, and IJ_Rhizo-the former being a commercial software and the latter being an open-source, freeware. We used simulated root images of known measurements (http://doi.org/10.5281/zenodo.1159845; Rose and Lobet 2018) which were previously used in Rose and Lobet (2019). A Root System Markup Language file had been previously generated using the root model ArchiSimple (Pagès *et al*. 2014) for 50 taproot and 50 fibrous root systems which were then stored as black and white JPEG image files (resolution of 1200 dpi) (Rose and Lobet 2019). Images were analyzed with a thresholding value set to 144 for the three software. Values for length, average diameter, surface area, and volume for the root images for IJ_Rhizo and WinRhizo™ were used from Rose and Lobet (2018) and batch analyzed to get the corresponding measurements for RVE.

To compare the efficacy of RVE, scanned grayscale root images of maize (*Zea mays* L.), wheat (*Triticum aestivum* L.), and various herbaceous and tree species were analyzed and compared with the image analysis results from WinRhizo™. The herbaceous species (Fig 4D) include 9 species sampled at ca. 115 days old: (*Bromus erectus, Dactylis glomerata, Holcus lanatus, Plantago lanceolata, Sanguisorba minor, Taraxacum officinale, Lotus corniculatus, Medicago sativa, Trifolium repens*) (Freschet *et al*. 2018). The maize (*Zea mays* L.) roots included scans (Fig 4E) from 40-day old plants across 30 cm depths for several nodes of roots (Guo and York, 2019). The wheat *(Triticum aestivum* L.) roots are entire root systems from 10-day old wheat seedlings grown in hydroponics (Guo *et al*. 2020). The tree roots are from the unpublished work of M. Luke McCormack and include root branches from 14 gymnosperm species sampled from the field *(Cephalotaxus harringtonii, Ephedra distachya, Chamaecyparis pisifera, Ginkgo biloba, Juniperus chinensis, Larix decidua, Metasequoia glyptostroboides, Picea abies, Pinus resinosa, Pinus strobus, Sciadopitys verticillata, Taxus cuspidata, Taxodium distichum, Tsuga canadensis*). All these image sets were scanned separately in different labs using Epson scanners with transparency units at resolutions of 600 DPI for maize, wheat, and herbaceous species, and 800 DPI for the tree species. Measurements were extracted for length, average diameter, surface area, and volume. In RhizoVision Explorer, the images were analyzed with the following analysis options: (i) *Analysis mode* as ‘Broken roots’, (b) *Image thresholding level* of 205, (c) *Filter non-root objects* as 8 mm^2^, (d) *Filter holes in root objects* set to false, (e) *Enable edge smoothing* set to false, and (f) *Root pruning threshold* set to 5. In the case of WinRhizo™, parameters were kept as the original used by the respective image providers, thresholding values were set as 205 in the manual mode for maize and wheat roots, automatic for herbaceous, and global Lagarde with a 64-pixel region size for the tree roots. In both software, the respective resolution was used. Fig 4 shows the input images including a root crown image as well as the corresponding processed images highlighting the skeletal structure of the roots that are colored based on various diameter ranges. These 4 image sets of roots from several plant species are available in a public repository and can be downloaded at http://doi.org/10.5281/zenodo.4677751 (Dhakal *et al*. 2021b).

### Statistical analysis

Statistical analyses were performed using R (R Core Team 2020) through RStudio (RStudio Team 2020). The R package “ggplot2” (Wickham 2016) was used for data visualization. Other packages used in the analysis were “ggpmisc” (Pedro 2020), “tidyverse” (Wickham *et al*. 2019), “ggpubr”, “magrittr”, “readxl”, and “Metrics” (Hamner and Frasco 2018). The R code and tabular data used in this study are available in a public repository and can be downloaded at http://doi.org/10.5281/zenodo.4677553 (Dhakal *et al*. 2021c).

Linear regression was used to compare paired measurements for ground truth data and estimated measurements. Slope (α), intercept (β), and coefficient of determination (R^2^) of the linear regression between the ground truth and the estimated values were calculated. In order to quantify the prediction error for the software, various performance metrics such as mean bias error (MBE), root mean square error (RMSE), and determination coefficient (R^2^) were used. The R package ‘Metrics’ was used for calculating the prediction errors. RMSE is the measure of the standard deviation of the residuals. MBE is an index to quantify whether the estimates are under- or overestimated. A good prediction results in the narrow spread of the residuals. Likewise, MBE also informs the direction and magnitude of the bias.

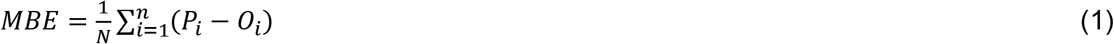

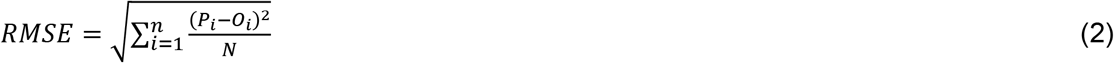

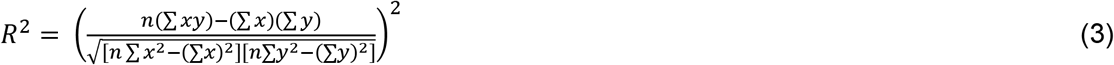

where P_i_ is the estimated value, Oi is the measured value, and N is the sample size. Lower values for both MBE and RMSE indicate a more accurate prediction.

## Results

### Validation with copper wires

#### Length

All IJ_Rhizo, RVE, and WinRhizo™ generated accurate length data for the copper wires. Determination coefficients were 1 for all software (*p* < 0.001), indicating a strong correlation between the ground truth measurements and the estimated length (Fig 5A). RMSE for estimated length for scanned copper wire images for IJ_Rhizo, RVE, and WinRhizo™ was 10.74 mm, 4.56 mm, and 6.67 mm, respectively. Results show that IJ_Rhizo overestimated length (MBE= 7.84 mm), RVE slightly overestimated length (MBE = 0.02 mm), and WinRhizo™ slightly underestimated the length (−4.2 mm) (Table 2).

**Figure 5.**
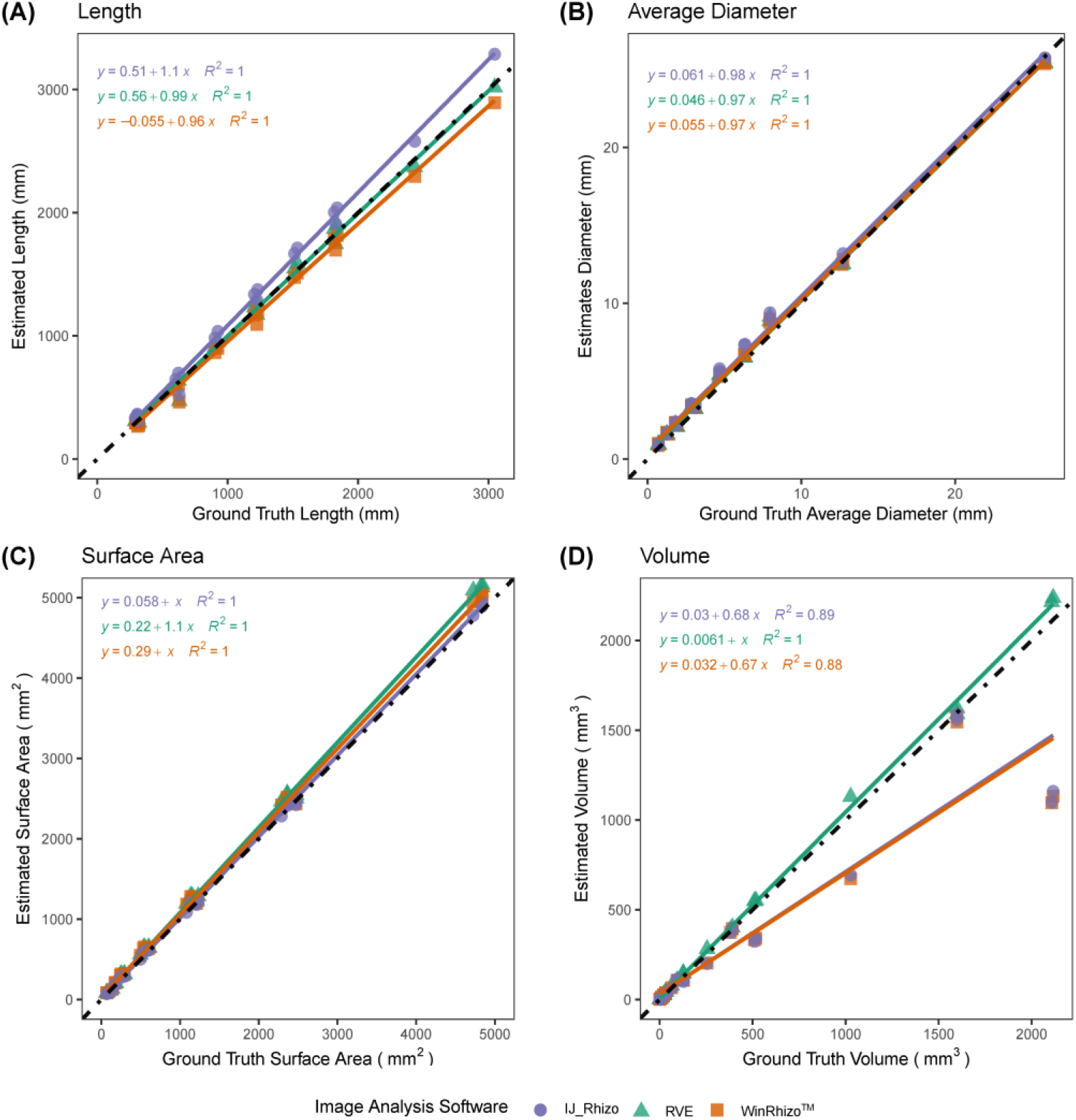
Comparison of physical measurements (length, average diameter, surface area, and volume) of various gauges of copper wires to the trait estimates generated from scanned images acquired at 600 dpi and analyzed using IJ_Rhizo, RhizoVision Explorer (RVE), and WinRhizo™, at a threshold value of 191. The fitted regression line for RVE and WinRhizo™ are shown in green and orange, respectively. The dotted black line represents a 1:1 relationship. A, B, C, D show length, average diameter, surface area, and volume, respectively.

**Table 2.** Comparison of various traits measured by IJ_Rhizo, RhizoVision Explorer (RVE), and WinRhizo™ using scanned copper wires images (n = 54) of known measurements. Length and average diameter are given in millimeters, and surface area and volume are given in squared millimeters and cubic millimeters, respectively.

| Traits | Root Mean Square Error | Mean Bias Error | Coefficient of Determination | p-value | Software |
| --- | --- | --- | --- | --- | --- |
| Length (mm) | 10.74 | 7.84 | 1 | <.001 | IJ_Rhizo |
|  | 4.56 | 0.02 | 1 | <.001 | RVE |
|  | 6.67 | -4.2 | 1 | <.001 | WinRhizo™ |
| Avg. Diameter (mm) | 0.06 | 0.05 | 1 | <.001 | IJ_Rhizo |
|  | 0.05 | 0.03 | 1 | <.001 | RVE |
|  | 0.05 | 0.04 | 1 | <.001 | WinRhizo™ |
| Surface Area (mm<sup>2</sup>) | 0.4 | 0.2 | 1 | <.001 | IJ_Rhizo |
|  | 1.42 | 1.03 | 1 | <.001 | RVE |
|  | 1 | 0.72 | 1 | <.001 | WinRhizo™ |
| Volume (mm<sup>3</sup>) | 0.28 | -0.1 | 0.89 | <.001 | IJ_Rhizo |
|  | 0.04 | 0.02 | 1 | <.001 | RVE |
|  | 0.28 | -0.1 | 0.88 | <.001 | WinRhizo™ |

#### Average Diameter

For average diameter, all three root image analysis software (IJ_Rhizo, RVE, and WinRhizo™) returned nearly perfect fits (R^2^ =1; *p* < 0.001) with similar RMSE values (Fig 5B). IJ_Rhizo, RVE, and WinRhizo™, all slightly overestimated average diameter values (MBE 0.03–0.05 mm, respectively). RVE had the lowest MBE (0.03 mm) among the three software (Table 2).

#### Surface Area

The surface area for the scanned copper wire images was accurately estimated with all three software IJ_Rhizo, RVE, and WinRhizo™ (R^2^= 1) and RMSE values of 0.40 mm^2^, 1.42 mm^2^, and 1mm^2^, respectively (Fig 5C). All of these software overestimated surface area (MBE 0.2 mm^2^, 1.03 mm^2^, and 0.72 mm^2^, respectively) (Table 2).

#### Volume

RVE had a strong agreement between the ground truth measurements and estimated volume from the scanned copper wire images (R^2^=1, p<0.001). RMSE for RVE volume estimates was 0.04 mm^3^ with slight volume overestimation (MBE = 0.02 mm^3^). Both IJ_Rhizo and WinRhizo™ had a substantially lower agreement between the measured and estimated volume (R^2^ of 0.88 and 0.89, respectively; p <0.001) with an RMSE value of 0.28 mm^3^. The MBE value for both WinRhizo™ and IJ_Rhizo is −0.1 mm^3^, a slight underestimation (MBE =−0.1 mm^3^), as supported by the slope of the regression line 0.67 (Fig 5D, Table 2).

### Performance of IJ_Rhizo, RVE, and WinRhizo™ using simulated root systems

The determination coefficients obtained for the regression between estimated length and ground truth values for IJ_Rhizo, RVE, and WinRhizo™ indicated a strong agreement between the estimated length and ground truth measurements (R^2^ = 0.99, *p* < 0.001) (Fig 6A).

**Figure 6.**
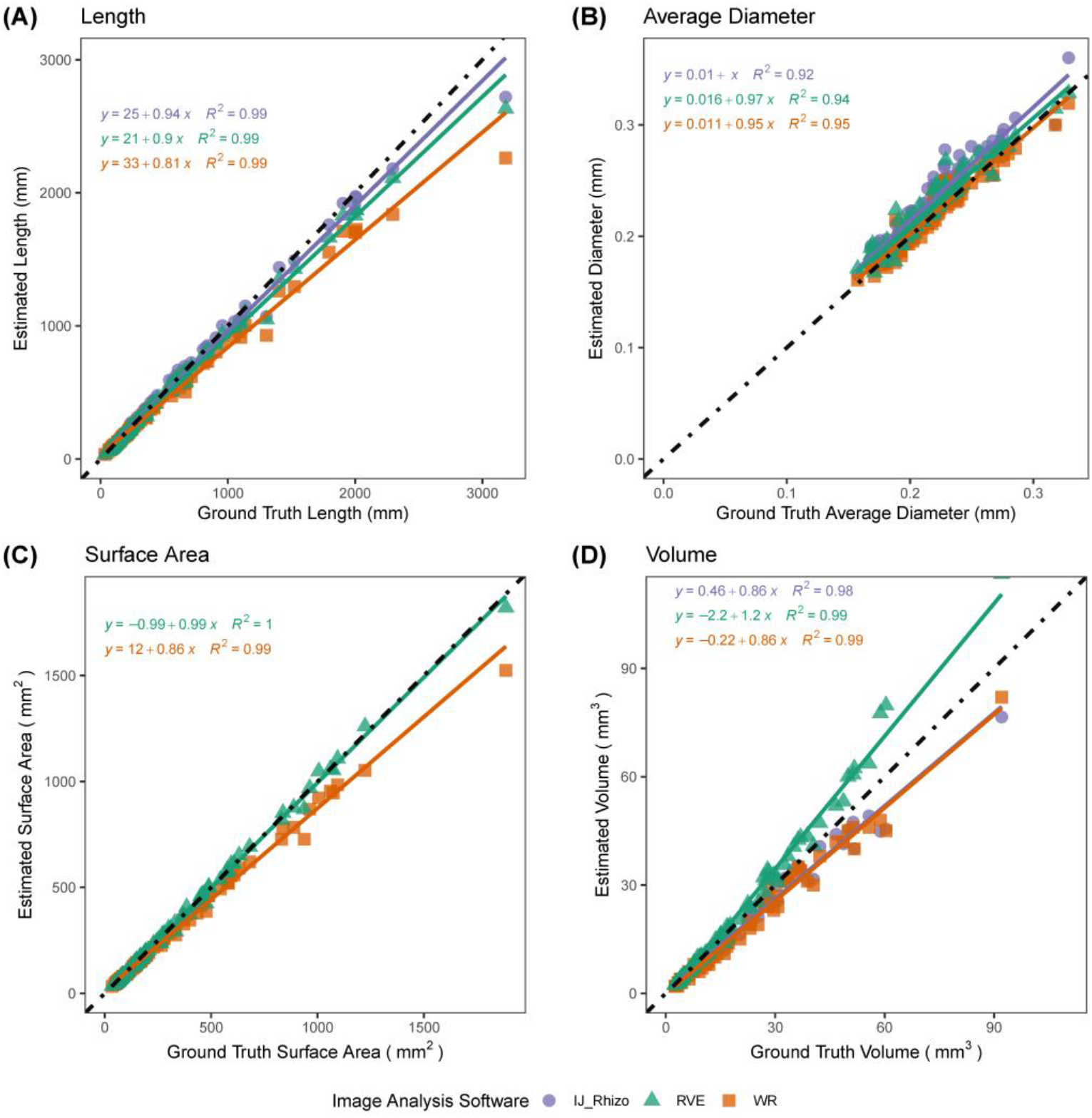
Comparison among IJ_Rhizo, RhizoVision Explorer, and WinRhizo™ for various trait estimates (n =294) against ground truth measurements obtained from simulated root images. For each trait, a linear regression model is fitted for IJ_Rhizo, RVE, and WinRhizo™ as shown in purple, orange, and green, respectively. The dotted black line represents a 1:1 relationship. A, B, C, D show length, average diameter, surface area, and volume, respectively.

IJ_Rhizo had the lowest RMSE (58.18 mm) and MBE (−5.46) values, followed by RVE (RMSE = 74.86 mm, MBE = −29.14 mm) and WinRhizo™ (RMSE = 137.72 mm, MBE = −64.50 mm), respectively (Table 3). The MBEs are positive indicating a tendency to underestimate length for the three software.

**Table 3.** Comparison of various traits measured by IJ_Rhizo, RhizoVision Explorer (RVE), and WinRhizo™ using simulated root images of dicot and monocot root systems (n = 294) of known measurements. Length and average diameter are given in millimeters, and surface area and volume are given in squared millimeters and cubic millimeters, respectively.

| Traits | Root Mean Square Error | Mean Bias Error | Coefficient of Determination | p-value | Software |
| --- | --- | --- | --- | --- | --- |
| Length (mm) | 58.18 | -5.46 | 0.99 | <.001 | IJ_Rhizo |
|  | 74.86 | -29.14 | 0.99 | <.001 | RVE |
|  | 137.72 | -64.50 | 0.99 | <.001 | WinRhizo™ |
| Avg. Diameter (mm) | 0.02 | 0.01 | 0.92 | <.001 | IJ_Rhizo |
|  | 0.01 | 0.01 | 0.94 | <.001 | RVE |
|  | 0.01 | 0.01 | 0.95 | <.001 | WinRhizo™ |
| Surface Area (mm<sup>2</sup>) | NA | NA | NA | NA | IJ_Rhizo |
|  | 15.19 | -2.59 | 1 | <.001 | RVE |
|  | 58.86 | -31.91 | 0.99 | <.001 | WinRhizo™ |
| Volume (mm<sup>3</sup>) | 3.68 | -5.46 | 0.98 | <.001 | IJ_Rhizo |
|  | 4.58 | 1.79 | 0.99 | <.001 | RVE |
|  | 3.93 | -2.71 | 0.99 | <.001 | WinRhizo™ |

Likewise, the determination coefficients obtained for the regression between estimated average diameter and ground truth average diameter values for IJ_Rhizo, RVE, and WinRhizo™ indicated a strong agreement between the estimated length and ground truth measurements (R^2^ > 0.9, *p* < 0.001) (Fig 6B). RVE and WinRhizo™ had lower RMSE values (0.02 mm) compared to IJ_Rhizo. A negative MBE (0.01) indicated a tendency to overestimate diameter for the three software slightly (Table 3).

For surface area, because Rose and Lobet (2019) did not include the analysis, the data were not presented for IJ_Rhizo. Determination coefficients were high for both RVE and WinRhizo™ (R^2^>0.99, p <0.001) (Fig 6C). WinRhizo™, however, returned the highest RMSE (58.86 mm^2^) and MBE values (−31.91 mm) compared to RVE (RMSE = 15.19 mm and MBE = −2.59 mm) (Table 3).

Linear regression analysis for volume confirms a strong correlation between the ground truth measurement and the estimated volume for the three software (R^2^> 0.98, *p* <0.001) (Fig 6D). Although, the RMSE values were highest for RVE (4.58 mm^3^), it had the lowest MBE (1.79 mm^3^) compared to IJ_Rhizo (RMSE = 3.68 mm^3^, MBE = −5.46 mm^3^) and WinRhizo™ (RMSE = 3.93 mm^3^, MBE = −2.71 mm^3^) (Table 3).

### Comparisons across roots of various plant species

For length, average diameter, surface area, and volume, WinRhizo™ trait values were typically underestimated relative to RhizoVision Explorer across all the species based on intercept and slope estimates from linear regression (Fig 7). However, typical R^2^ values greater than 0.9 indicate that the relative ranking of individual data points was relatively consistent.

**Figure 7.**
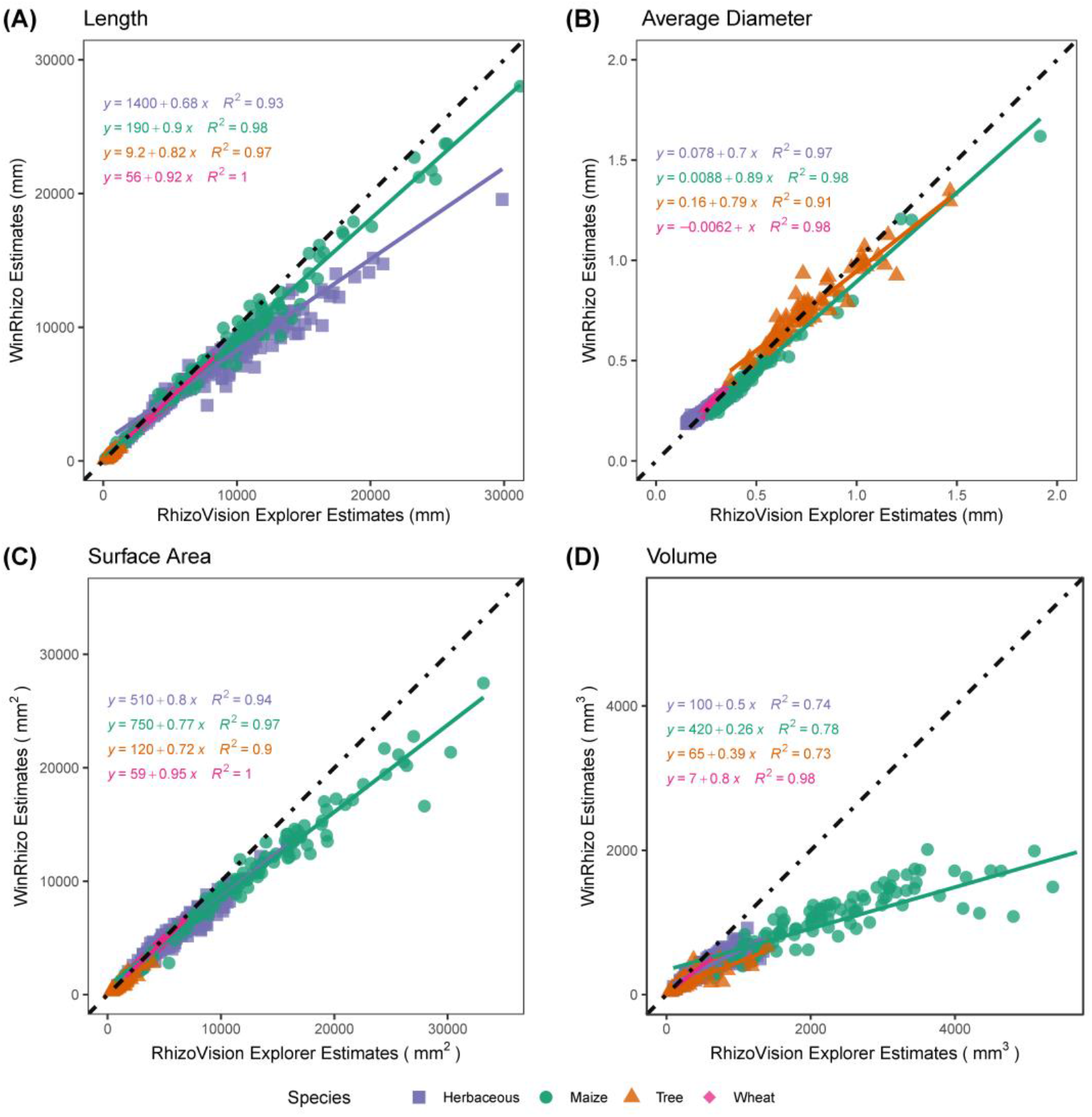
Comparison between RhizoVision Explorer and WinRhizo™ for various trait estimates across 4 image sets representing diverse plant species. For each trait, a linear regression model is fit for herbaceous species, maize, tree species, and wheat as shown in purple, green, orange, and pink, respectively. The dotted black line represents a 1:1 relationship. A, B, C, D show length, average diameter, surface area, and volume, respectively.

## Discussion

In this paper we have introduced RhizoVision Explorer (RVE), a freely available, easy-to-use, GUI-based, precompiled executable software for personal computer (PC) users. To facilitate adoption, the open-source software can be executed easily on most Windows PCs (Windows 8.1 or 10) without the need for dedicated GPUs or cloud-hosted services. The program is developed for a modular approach with the implementation of open-source libraries such as *Qt, OpenCV*, and a plugin system. Developers can easily build upon the existing open-source software codebase to fit their root research pipeline and share the software with the root phenotyping community. The software provides interactive and batch processing modes, region-of-interests functionality, and numerous options for segmentation, filtering, feature extraction, and output display.

To validate the physical measures provided by the software, a novel image set was created using six diameters of copper wires with known lengths imaged in 28 combinations and scanned with a similar protocol as for plant roots. RhizoVision Explorer showed excellent agreement with the ground truth values for total length, average diameter, surface area, and volume. In some cases, reflections or glare from the smooth copper wire produced non-ideal images with minor imperfections. A published image and dataset were used to compare RVE with previous results obtained for IJ_Rhizo and WinRhizo™. In general, the three software performed similarly and their inaccuracies can largely be attributed to the ground truth data was based on simulated 3D root systems, but that image analysis results are based on single perspective 2D projections that increase root occlusion and overlap. When the original 3D images are flattened, many of the fine roots are effectively combined into larger diameter structures while simultaneously decreasing root length, as seen in Fig 6A showing underestimation of length by all software. Larger diameter structures would tend to inflate volume estimates, derived from the squared radius in a cylinder formula, while decreased length would tend to decrease the volume. Therefore, RhizoVision Explorer may overestimate the volume (Fig 6D) due to the increased apparent diameters observable in the crown of the simulated root images (see green medial axis visible in the upper right of Fig 4H), while WinRhizo and IJ_Rhizo underestimate volume because of their bias in using the average diameter and length as discussed below. In practice, this type of root occlusion may be minimized at the imaging stage by ensuring roots are spread with minimal overlap. The root image analysis community would benefit from further ground truth datasets that correct some of these issues, such as 2D simulated roots of both connected and unconnected types, with various overlap, root hair presence, distortion, and diameter heterogeneity.

For performance evaluation of RhizoVision Explorer with typical scanned root images, and to aid future root algorithm development, another image set containing wheat, maize, wild herbaceous species, and tree species was created. In this case, only a comparison with the commonly used WinRhizo™ was conducted. Overall, the results show good agreement between the two software, except WinRhizo™ substantially underestimated root volume relative to RhizoVision Explorer. Previous work had indicated that WinRhizo™ and IJ_Rhizo inaccurately calculate volume based on the average diameter of all roots in an image and the total length of all roots in the image (Delory *et al*. 2017; Rose and Lobet 2019). This method assumes all roots are of the same (average) diameter, but diameters of roots within a sample can vary 10-fold. In such samples with a great variation of diameter, the average diameter will be biased by the smaller diameter roots and the volume will be underestimated. RhizoVision Explorer does not suffer from this inaccurate method, because surface area and volume are calculated by assuming that every pixel in the skeleton is a cylinder and summing across the image or diameter range. The severe volume underestimation possible when using WinRhizo™ or IJ_Rhizo is readily observable in the copper wire image set (Fig 4B,G), where RhizoVision Explorer gave the correct result, but WinRhizo™ or IJ_Rhizo only reported half of the volume in the most extreme cases (Fig 5D). When comparing results between WinRhizo™ and RVE across the image sets of various crop and wild species, reasonable agreement is seen for length, average diameter, and surface area, but in some maize root images with significant diameter heterogeneity between nodal and lateral root diameters (see Fig 4E,J), the WinRhizo™ volume estimate was five-fold less (Fig 7D). Volume is an important measurement, typically used as a proxy for root mass and also for the calculation of root tissue density. These measures relying on volume that exist in the literature and in public root databases like FRED (Iversen *et al*. 2017) or GRoot (Guerrero-Ramírez *et al*. 2021) should be carefully evaluated before use.

RhizoVision Explorer uses global grayscale thresholding to encourage imaging platforms that maximize the contrast of roots with the background, such as by using the RhizoVision Crown platform for root crown imaging using a backlight (Seethepalli *et al*. 2020), or use of a flatbed scanner with a transparency unit (or top light). Therefore, many types of complex images produced with blue paper screens, rhizoboxes, or minirhizotrons may not work directly, which presents a limitation of the software. Image analysis tools have built-in assumptions of the input image on which the analysis operations are based, such as high contrast between the roots and the background. Users of these tools should carefully consider each tool before collecting or processing image data to ensure accurate and representative results. Image preprocessing steps such as cropping, filtering, or thresholding are often required separately before analysis in software such as ImageJ. However, several software tools have become available over the past few years for machine learning approaches for root segmentation from complex backgrounds, with RootPainter being a GUI-based example (Smith *et al*. 2020). In addition, ImageJ has many thresholding options, including color, which could be applied in batch processing. Segmented images can be produced in such a companion software, and then processed in RhizoVision Explorer for feature extraction.

Future development of RhizoVision Explorer will focus on correcting any bugs identified, improving the user interface for convenience, and expanding features measured, especially for differentiating root classes such as laterals and axial roots or branch orders. In addition, the presence of root hairs in images can drastically increase all measures but this issue has rarely been discussed in the literature. Methods to address problems from root hairs may include blurring routines before image segmentation, or additional methods to prune the skeleton. In order to generate a skeleton that encodes a more accurate topology, in the future further refinements to define branch points and delete invalid root tips will need to accommodate the false loops as seen in Fig 3. Most topological analysis is based on simple considerations of the skeleton branching, however other considerations such as child and parent root diameters may help to refine the correct topology. As improvements are made in identifying root type or order, more measurements at the level of individual roots may be beneficial such as lateral root insertion angles or diameters. The software is open-source and community development is highly encouraged. RhizoVision Explorer is intended for roots, however, it is suitable for measuring leaf area from scanned images, and the plugin-based software architecture could be extended for other phenotyping needs.

## Conclusion

RhizoVision Explorer is an image analysis tool that is intended to enable many more researchers to be able to routinely measure plant roots to answer a variety of biological questions. The graphical user interface and ready-to-run download for Windows, as seen in other root-related software (Pound et al. 2013), lowers barriers and increases accessibility to image analysis. The underlying software architecture is modular and with plugin support that is suitable for modifying for other phenotyping, or other image analysis needs. Future improvements include greater topology analysis with the ability to predict root order, to provide more features at the diameter bin or order level such as average diameter and angle. In summary, this open-source software builds on a range of previous software to propose a user-friendly, generalist, collectively improvable, and all-inclusive tool that will facilitate the standardization of root architectural and morphological measures.

## Data Availability

RhizoVision Explorer is available as a ready-to-run executable for Windows at https://doi.org/10.5281/zenodo.3747697 (Seethepalli and York 2020).

The open-source code for RhizoVision Explorer written in C++ is available at https://github.com/noble-research-institute/RhizoVisionExplorer on GitHub.

The copper wire image set is available in a public repository and can be downloaded at http://doi.org/10.5281/zenodo.4677546 (Dhakal *et al*. 2021a).

These 4 image sets of roots from several plant species are available in a public repository and can be downloaded at http://doi.org/10.5281/zenodo.4677751 (Dhakal *et al*. 2021b).

The statistical R code and tabular data used in this study are available in a public repository and can be downloaded at http://doi.org/10.5281/zenodo.4677553 (Dhakal *et al*. 2021c).

## Acknowledgments

The authors acknowledge nearly 30 beta testers not listed here that provided helpful feedback to improve the software before the public release, including in a survey answered by 20 testers. Especially, the authors thank Arthur Villordon both for his suggestions and his promotion of the software in the root crop community. The authors thank M. Luke McCormack for providing the tree root images.

## Contributions by the Authors

The software was conceptualized and developed by AS and LMY, and programmed by AS. KD acquired the wire image set, performed statistical analysis with contributions from AS, and generated all data plots and data tables. AS and LMY generated other figures. All authors gave feedback on the software and data analysis during development, contributed to writing the manuscript, and approved the final version for submission.

---

**Supplemental Table 1.** Combinations of 12 wires used for the copper wire image data set, ranging from 1 to 12 wires in a scan. For each of 6 wire gauges (diameters), two wires of approximately 30 cm length were used and denoted as wire 1 or 2. File names for the scans indicate combinations of the gauges and wire numbers used, and the number (#) of wires in a scan are provided.

| AWG Gauge | File Name | # |
| --- | --- | --- |
| 40 | G_40_W1_600_dpi.tif | 1 |
| 40 | G_40_W2_600_dpi.tif | 1 |
| 32 | G_32_W1_600_dpi.tif | 1 |
| 32 | G_32_W2_600_dpi.tif | 1 |
| 28 | G_28_W1_600_dpi.tif | 1 |
| 28 | G_28_W2_600_dpi.tif | 1 |
| 22 | G_22_W1_600_dpi.tif | 1 |
| 22 | G_22_W2_600_dpi.tif | 1 |
| 16 | G_16_W1_600_dpi.tif | 1 |
| 16 | G_16_W2_600_dpi.tif | 1 |
| 10 | G_10_W1_600_dpi.tif | 1 |
| 10 | G_30_W2_600_dpi.tif | 1 |
| 40 | MD_40_40_W12_600_dpi.tif | 2 |
| 40, 32 | Mixed_40_32_W1_600_dpi.tif | 2 |
| 40, 32 | Mixed_40_32_W2_600_dpi.tif | 2 |
| 40, 32 | MD_40_32_40_32_W1122_600_dpi.tif | 4 |
| 40, 32, 28 | Mixed_40_32_28_W1_600_dpi.tif | 3 |
| 40, 32, 28 | Mixed_40_32_28_W2_600_dpi.tif | 3 |
| 40, 32, 28 | MD_40_32_28_40_32_28_W111222_600_dpi.tif | 6 |
| 40, 32, 28, 22 | Mixed_40_32_28_22_W1_600_dpi.tif | 4 |
| 40, 32, 28, 22 | Mixed_40_32_28_22_W2_600_dpi.tif | 4 |
| 40, 32, 28, 22 | MD_40_32_28_22_40_32_28_22_W11112222_600_dpi.tif | 8 |
| 40, 32, 28, 22, 16 | Mixed_40_32_28_22_16_W1_600_dpi.tif | 5 |
| 40, 32, 28, 22, 16 | Mixed_40_32_28_22_16_W2_600_dpi.tif | 5 |
| 40, 32, 28, 22, 16 | MD_40_32_28_22_16_40_32_28_22_16_W1111122222_600_dpi.tif | 10 |
| 40, 32, 28, 22, 16, 10 | Mixed_40_32_28_22_16_10_W1_600_dpi.tif | 6 |
| 40, 32, 28, 22, 16, 10 | Mixed_40_32_28_22_16_10_W2_600_dpi.tif | 6 |
| 40, 32, 28, 22, 16, 10 | MD_40_32_28_22_16_10_40_32_28_22_16_10_W111111222222_600_dpi.tif | 12 |

